# CUAD biotechnology offers 0.5 USD per hectare for aphid control: innovative platform for low-cost plant protection

**DOI:** 10.1101/2024.05.03.592334

**Authors:** Volodymyr V. Oberemok, Yelizaveta V. Puzanova, Nikita V. Gal’chinsky

## Abstract

20 years ago it was difficult to imagine the use of nucleic acids in plant protection as insecticides, but today it is a reality. At the very beginning, new technologies often work inefficiently and are expensive; in the process of their development, qualitative changes occur that make new technologies accessible and work flawlessly. Invented in 2008, oligonucleotide insecticides based on CUAD (contact unmodified antisense DNA) platform have been improved and today possess characteristics that amaze the imagination: low carbon footprint, high safety for non-target organisms, rapid biodegradability in ecosystems, avoidance of target-site resistance. This next generation class of insecticides creates opportunities to develop insecticides which are well-tailored for a particular population of insect pest. CUAD biotechnology combines the achievements of molecular genetics, bioinformatics, and *in vitro* nucleic acid synthesis providing a simple and flexible platform for plant protection. Aphids, as one of the key pests of important agricultural crops that shape food security, can be controlled by oligonucleotide insecticides at affordable price already today, ensuring effective control with minimal risks for the environment. In this article, low-dose concentration (0.1 ng/µL; 20 mg per hectare in 200 L of water solution) of 11-nt long oligonucleotide insecticide Schip-11 shows effectiveness on aphid *Schizolachnus pineti* causing 76.06 ± 7.68 mortality rate on the 12th day (p<0.05). At a consumption rate of 200 L per hectare, the price of the required amount of oligonucleotide insecticide will be about 0.5 USD/ha using liquid-phase DNA synthesis and makes oligonucleotide insecticides very competitive on the market. Also we show that non-canonical base pairing G:T (T:G) between target pre-rRNA and olinscide is well tolerated in aphids. Thus, non-canonical base-pairing should be taken into consideration during design of olinscides not to harm non-target organisms.

## 1. Introduction

To date, contact unmodified antisense DNA (CUAD) biotechnology is the only platform that uses short unmodified antisense DNA as contact oligonucleotide insecticides (briefly, olinscides, or DNA insecticides) [1,2,3]. DNA, as a programming molecule and polymer of natural origin, has always attracted researchers, but for a long time it was believed that unmodified oligonucleotides are very quickly degraded in all eukaryotic cells under the action of nucleases [4], including insects [5]. Practical application of the principles of antisense oligonucleotides (ASOs) was first formulated in Novosibirsk (Russia) by N. Grineva in 1967 [6]. In 1977, using this antisense approach to modify valine tRNA, N. Grineva and colleagues demonstrated that the method allows alkylation with a reagent bound to the corresponding oligonucleotide at certain points along the valine tRNA [7]. P. Zamecnik and M. Stephenson in 1978 were the first to use modified antisense DNA against Rous sarcoma virus in chicken embryo fibroblasts in a sequence-specific manner [8]. The development of antisense technologies has long been mainly towards medicine using modified antisense oligonucleotides [9]. After 20 years of research with modified antisense oligonucleotides, in 1998, the FDA licensed the first drug, Vitravene (Fomivirsen), based on the 21-mer phosphorothioate oligonucleotide [10]. This area of medicine continues to progress and has already been marked by the registration of other important drugs [11].

An unexpected turn with antisense oligonucleotides for plant protection came in 2008, when it was shown that short unmodified antisense DNA has significant insecticidal effect on insect pests [12]. For the first time, the equal sign between antisense oligodeoxyribonucleotides and contact insecticides was put in the experiments with spongy moth *Lymantria dispar*, which led to the birth of CUAD platform [13-16]. CUAD biotechnology has a number of features that distinguish it from all modern classes of chemical insecticides and plant protection technologies developing today: unmodified antisense DNA as active substance, DNA containment as mechanism of action, insect pre-rRNA and rRNA as target [1,2,17].

The CUAD platform works best on hemiterans from suborder Sternorrhyncha [3,17]. In the last few years CUAD biotechnology based on oligonucleotide insecticides (briefly, olinscides, or DNA insecticides) has been established as a powerful method against soft scale insects, armored scale insects, psyllids, mealybugs, and aphids, opening new frontiers in plant protection based on contact application of deoxyribonucleic acid [3,17]. Oligonucleotide insecticides possess low carbon footprint, high safety for non-target organisms, rapid biodegradability in ecosystems, and avoidance of target-site resistance. The idea of oligonucleotide insecticides attracted the attention of scientists and experts in plant protection [18-22] and to a certain extent the affordability of such insecticides remained in question. Our research team decided to lower the price of olinscides and find a serious group of insect pests on which oligonucleotide insecticides at low concentrations could have a significant insecticidal effect. Aphids from subfamily Lachninae turned out very sensitive to low-dose concentrations of oligonucleotide insecticides.

Aphids (Hemiptera: Aphididae) are significant economic pests that are found globally. Aphids feed on phloem [23] and cause substantial economic losses mainly spreading plant viruses, and producing honeydew [24-26]. Among aphids around 100 species are considered to be agricultural pests of a wide range of crops. They are major insect pests of various plants, including alfalfa, wheat, potato sugar beet and tobacco [27]. The damage caused by aphids amounts to hundreds of millions of dollars a year [28]. Their management is challenging because the mobility of aphids is extremely high [29]. Also, these pests reproduce predominantly asexually [30], one female leaves 10-90 offspring in 7-10 days and therefore, theoretically, could produce billions of offspring in one growing season in the absence of mortality factors [31]. Chemical insecticides have been used to control aphids, and these pests quickly develop resistance to various classes of chemical insecticides, including neonicotinoids, carbamates, organophosphates, organochlorines, and pyrethroids [32]. This prompts the search for new insecticides with advanced characteristics and multi-decade utility.

In this article we use aphid *Schizolachnus pineti* for the experiments. *S. pineti* is a serious pest of *Pinus* spp., but especially on young Scots pine (*Pinus sylvestris*) where it forms dense colonies in rows along the previous year’s needles [33].

## 2. Results

### 2.1. Mortality of S. pineti after contact application of olinscide Schip-11

After treatment of *S. pineti* with olinscide Schip-11 in concentration 200 ng/μL mortality of the pest reached 24.32 ± 1.37 %, 61.03 ± 2.17 %, 76.56 ± 3.67 %, and 84.19 ± 3.84 % on the 1^st^, 2^nd^, 3^rd^, and 4^th^ day, respectively (Figure 1, A). Of, note the same olinscide (5’-TGT-GTT-CGT-TA-3’, Macsan-11) with perfect complementarity to target ITS2 of polycistronic rRNA transcript caused significant mortality of closely related species, chrysanthemum aphid *Macrosiphoniella sanborni*, in concentration 100 ng/µL. It led to 67.15 ± 3.32 % mortality rate of the chrysanthemum aphid after a single treatment and 97.38 ± 2.49 % mortality rate after a double treatment (with daily interval) on the 7^th^ day [34]. Thus, we see substantial potential of ITS2 region of polycistronic rRNA transcript as a target for olinscides. Moreover, ITS regions of rRNA genes are moderately variable (Figure 4) in comparison with regions of 18S, 5.8S, and 28S rRNAs [35] allowing the creation of unique sequences of oligonucleotide insecticides. The length of an oligonucleotide insecticide ∼ 11 nt makes it possible to create selective oligonucleotide insecticides with a uniqueness frequency equal to 1/4.19·10^6^ and is obviously enough to be used in most agrocenoses [36].

**Figure 1.**
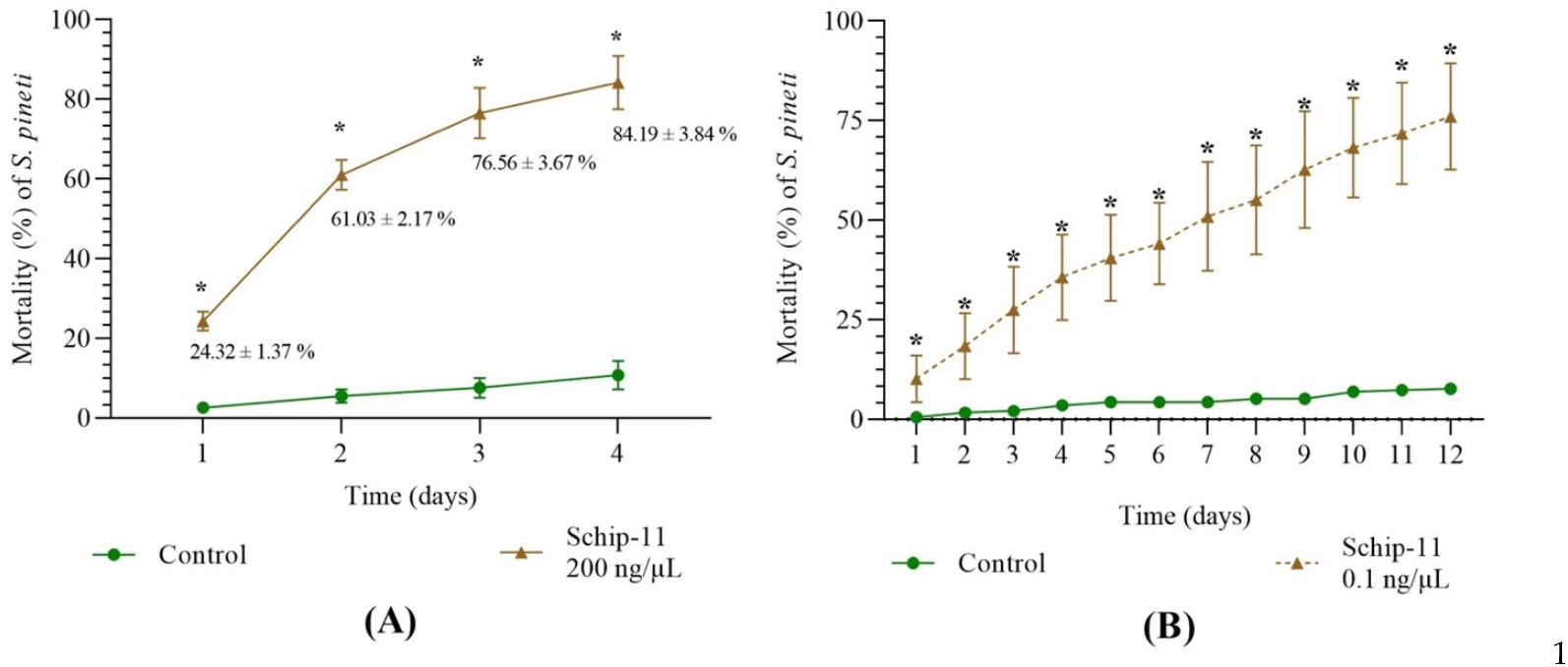
Dynamics of mortality of *S. pineti* after treatment with water and oligonucleotide insecticide Schip-11 in different concentrations: (A) 200 ng/µL; (B) 0.1 ng/µL; The significance of the difference between Schip-11 group and water-treated Control groups is indicated by * p<0.05.

The use of olinscides could solve, or at least improve, the fundamental problem of insecticide selectivity. The results of our previous work with aphid *M. sanborni* showed that the change of just one nucleotide at the 1^st^ (5′-end, T to A) and 11^th^ (3′-end, A to T) positions leads to dramatical decrease in biological efficiency of target 11-nucleotides long olinscide Macsan-11 [34]. At the same time on scale insects, *Dynaspidiotus britannicus* and *Aonidia lauri*, we showed that non-canonical base-pairing, such as A:? (?:A) and G:T (T:G) [37-39] may occur between olinscides and imperfect sites of rRNAs of non-target organisms. [1]. Here for the first time we show that non-canonical base pairing G:T (T:G) between target pre-rRNA and olinscide is also well tolerated in aphids. Definitely, non-canonical base-pairing should be taken into consideration during design of olinscides not to harm non-target organisms.

After treatment of *S. pineti* with olinscide Schip-11 in concentration 0.1 ng/μL mortality of the pest reached 10.18 ± 3.36 %, 18.44 ± 4.79 %, 29.51 ± 5.35 %, 35.67 ± 6.19 %, and 76.06 ± 7.68 % on the 1^st^, 2^nd^, 3^rd^, 4^th^, and 12^th^ day, respectively (Figure 1, B). Interestingly, graph of dynamics of mortality of the pest differs from the standard hyperbolic form which is characteristic for olinscides in concentration 100-200 ng/µL [1,36,40,41] and represents almost linear graph. It should also be noted that insect mortality occurs more slowly when concentration of olinscide is 0.1 ng/µL. Similar insect mortality (≈76%), obtained on 12^th^ day in the group with a concentration of 0.1 ng/µl, was achieved already on the 3^rd^ day in the group with a concentration of 200 ng/µl of olinscide.

Of note, closely related species of aphids from the same subfamily Lachninae, *Cinara pinea* and *Eulachnus rileyi*, also showed high sensitivity to Schip-11 in 0.1 ng/µL concentration. On the 12^th^ day, mortality of *C. pinea* and *E. rileyi* comprised 63.66 ± 19.81 % and 67.73 ± 9.16 %, respectively in comparison with water-treated Controls (9.14 ± 0.83 % and 8.31 ± 2.11 %, respectively) (p<0.05). It shows high reproducibility of results and perspective of using low-dose concentrations of olinscides.

### 2.2. Olinscide Macsan-11 significantly decreases concentration of polycistronic rRNA transcript of S. pineti (investigation of dead insects)

In this article we decided to investigate concentration of polycistronic rRNA transcript containing target ITS2 region for olinscide Schip-11 and used dead individuals. We found significantly decreased concentration of polycistronic rRNA transcript in dead insects treated with water solutions of olinscides in both concentrations, 200 ng/µL and 0.1 ng/µL, compared to water-treated Controls (p<0.05).

On the 1^st^ and 3^rd^ day, concentration of polycistronic rRNA was 15.62 and 9.09 times lower compared to water-treated Control for 200 ng/µL of Schip-11. For 0.1 ng/µL group, on the 1^st^ and 3^rd^ day, concentration of polycistronic rRNA was 17.85 and 45.45 times lower compared to water-treated Control (Figure 2).

**Figure 2.**
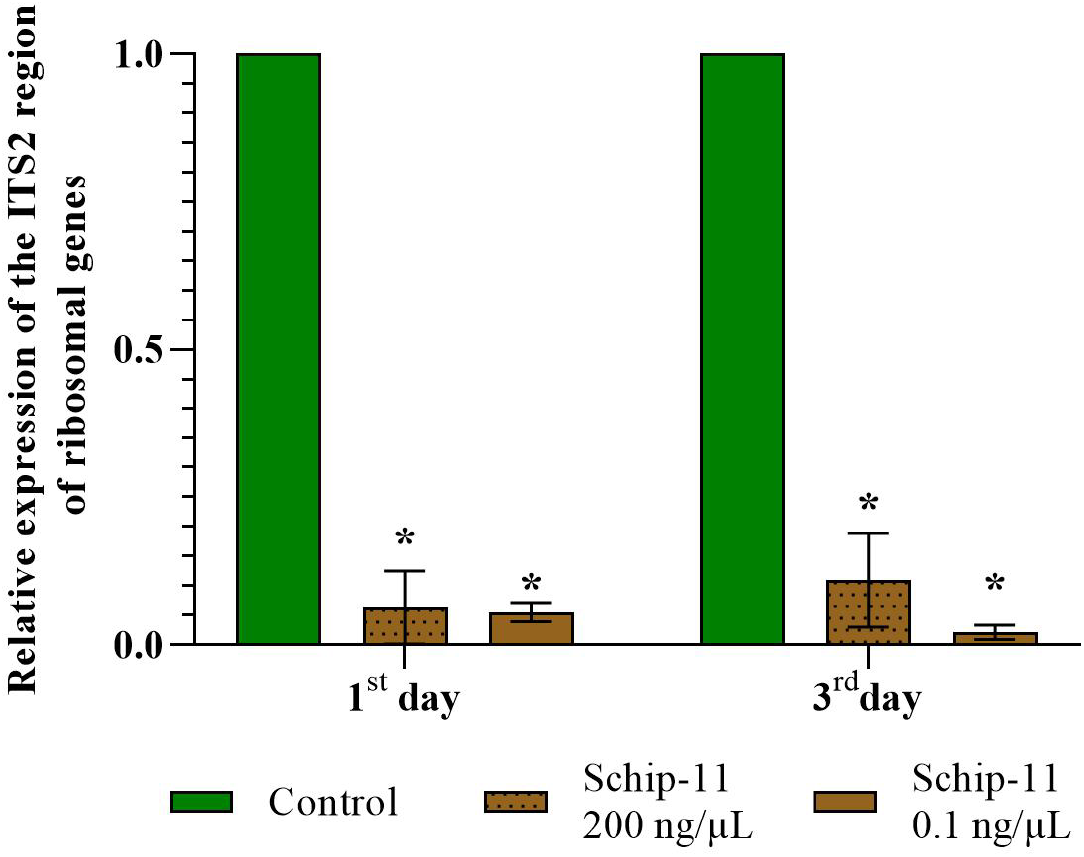
Relative concentration of polycistronic rRNA transcript of *S. pineti* after treatment with oligonucleotide insecticide Schip-11 in different concentrations (200 ng/µL; 0.1 ng/µL) on the 1^st^ and 3^rd^ day; water-treated Control was taken as 1 (100%). The significance of differences between Schip–11 group and water-treated Controls is indicated by * at p < 0.05.

Olinscides act on sternorrhynchans through DNA containment mechanism consisting of 2 steps: 1) rRNA and/or pre-rRNA ‘arrest’ and hypercompensation of target rRNA; 2) target rRNA and/or pre-rRNA degradation recruiting RNase H [1,2]. For alive individuals of chrysanthemum aphid *M. sanborni* we detected gradually decreasing hypercompensation of target rRNA in 1-3 days after treatment [34] while here we show that dead individuals have decreased concentration of target rRNA. Obviously, increased concentration of target rRNA is better than decreased one in comparison with water-treated Control and insects stay alive [1]. Nevertheless, after peak of target rRNA hypercompensation, we also see decrease in target rRNA concentration in alive individuals of insect pests caused by action of RNAse H [1,34,36].

## 3. Discussion

Low concentrations of oligonucleotide insecticides (0.1 ng/µl) showed high insecticidal potential against aphids. The detected high mortality rate indicates an effective and targeted effect of unmodified antisense DNA on the pest. It should be noted that at a comparable concentration (∼0.07 ng/µl) dsRNA also have a significant insecticidal effect on the Colorado potato beetle [42]. In parallel with oligonucleotide insecticides, a new class of insecticides based on dsRNA is being developed, the action of which is based on RNAi. RNA biocontrols show the best results on coleopterans [43], and much worse on hemipterans [44]. In this context, oligonucleotide insecticides and RNA biocontrols as 2 next-generation classes of insecticides, are able to complement each other’s action in complex preparations for wide range of pests from different orders, especially against those that have shown resistance to many different compounds from major insecticide classes [45].

Obtained results, show perspective of using olinscides against conifer aphids, including *S. pineti, E. rileyi, C. pinea*. Using cold fog generators and big cold fogging machines it is possible to treat vast territories of conifer forests without harm to natural enemies (wasps, mites, beetles, etc.) of insect pests and other non-target organisms. Aphids form an important part of many food chains and can be part of a healthy garden or forest ecosystem. Thus, olinscides can control a distinct pest species while closely related species will stay unharmed. Using unique complementary sequences to target pre-rRNAs and rRNAs of an insect pest it is possible to create well-tailored olinscides with minimal risks to balance of an ecosystem. As a molecule of natural origin, olinscides do not reduce biodiversity, do not impact soil health, and do not accumulated in ecosystems. We can say that olinscides degrade almost immediately recruiting ubiquitous DNases after they act on insect pests [1,16,34].

CUAD biotechnology, as well as double-stranded RNA technology, has achieved a significant reduction in the cost of nucleic acid synthesis using innovative methods for production of nucleic acids *in vitro* [46,47]. CUAD biotechnology has become significantly cheaper due to liquid phase synthesis [48]. One of market leaders in liquid phase synthesis, Sumitomo Chemical Co., Ltd. (Tokyo, Japan), offers the synthesis of 1 kg of un-modified oligonucleotides 11 nt long for 25,000 USD (personal communication). In the case of using non-optimized solid-phase DNA synthesis, which is available in many laboratories around the world, including ours, the cost of synthesizing 1 kg of unmodified oligonucleotides 11 nt long will be about 1 million USD. Thus, at a consumption rate of 200 L at a concentration of 0.1 mg/L (or 0.1 ng/µL) per hectare, the price of the required amount of oligonucleotide insecticide will be about 0.5 USD when using liquid-phase DNA synthesis. This price allows to increase the frequency of treatments with oligonucleotide insecticides in real conditions. If non-optimized solid-phase DNA synthesis is used, which is available in many laboratories around the world, including ours, the cost of synthesizing the required amount of oligonucleotide insecticide per hectare will be 20 USD. This price will be convenient for investigations in the lab.

The results obtained allow us to look at CUAD biotechnology as a platform capable to occupy a significant part of the insecticide market. Most of the insect pests against which CUAD biotechnology is effective today are representatives from the suborder Sternorrhyncha, which primarily live in the subtropics and tropics, and to a lesser extent, in the temperate zone [3]. Oligonucleotide insecticides can make a significant contribution to the protection of plants from pests of coffee, cocoa, citrus fruits, cereals, and other important groups of agricultural plants that ensure food security.

## 4. Materials and Methods

### 4.1. Origin of material

The experiments were carried out in laboratory conditions on shoots of *Pinus sylvestris* L. (Coníferae: Pinaceae) (Figure 3). Shoots of trees with *S. pineti* were randomized; 3 shoots were taken for each experimental group; there were approximately >100 aphids on each shoot. Experiments were conducted in triplicate.

**Figure 3.**
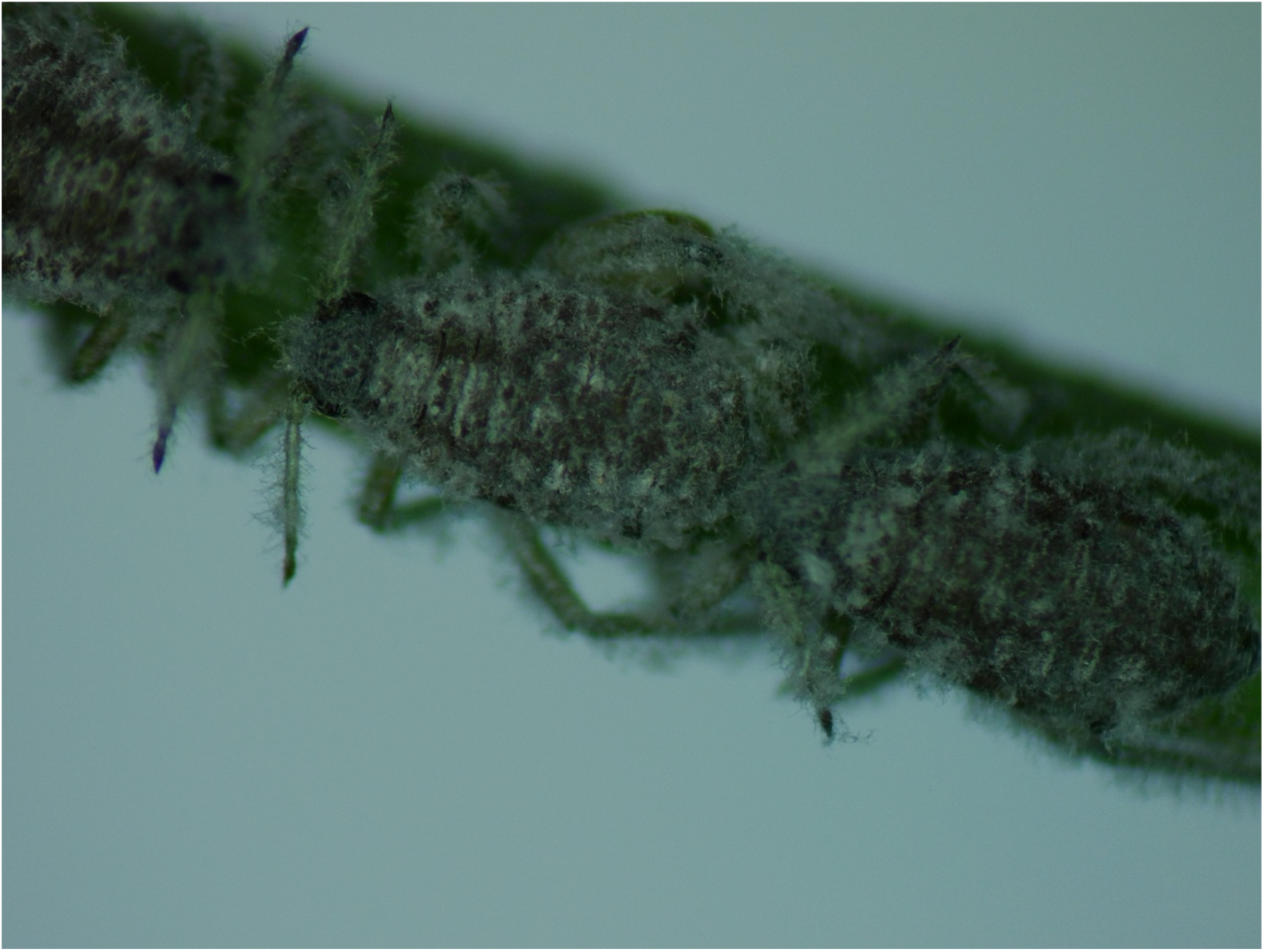
*S. pineti* on needle of *P. sylvestris*.

### 4.2. Design, synthesis, and application of oligonucleotide insecticide Schip-11

We designed oligonucleotide insecticide Schip-11 5’-TGT-GTT-CGT-TA-3’ which is almost complementary to the ITS2 region of pre-rRNA (polycistronic rRNA transcript) of *Schizolachnus pinet* (Figure 4). We decided to use Schip-11 sequence to find out if T:G (G:T) non-canonical base pairing in aphids is well tolerated as seen in scale insects [1].

**Figure 4.**
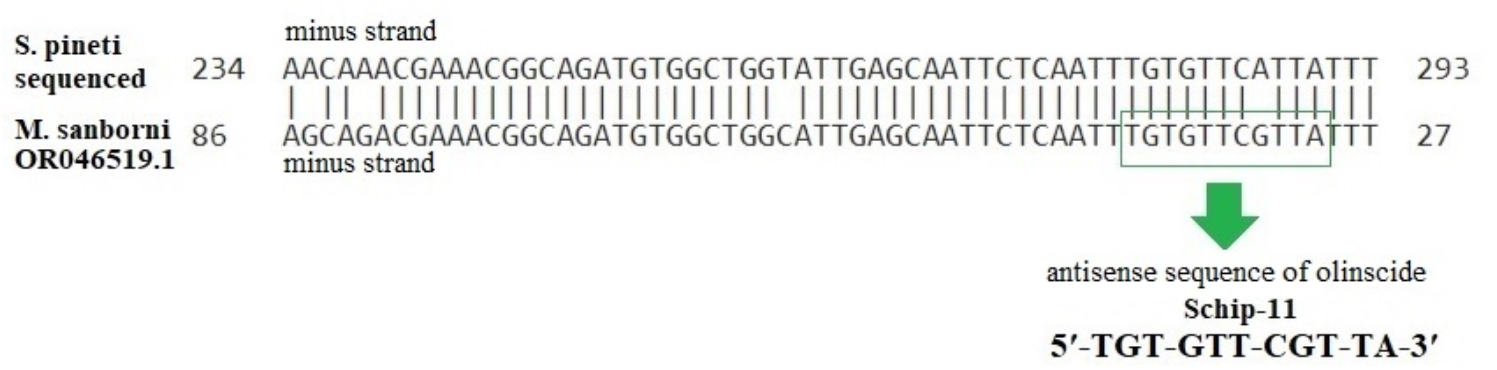
Alignment of the sequenced DNA fragment of *S. pineti* collected from nature and fragment of ITS2 region of rDNA of *M. sanborni* (GenBank: OR046519.1) performed using ClustalW 2.0.3.

The sequence of olinscide was synthesized using ASM 800E DNA synthesizer (BIOS-SET, Novosibirsk, Russia) according to standard phosphoramidite synthesis procedure. The synthesis was carried out in the direction from the 3′ to the 5′ end. After completion of all cycles of synthesis, the target oligonucleotide construct was removed from the solid-phase support; the removal of the protective groups was carried out overnight at 55 °C in a concentrated ammonia solution (analytical grade, “Vekton”, Saint Petersburg, Russia). Purification of the synthesized olinscide Schip-11 was performed on cartridges for purification of oligonucleotides OPS-12 (BIOSSET, Novosibirsk, Russia). A BactoSCREEN analyzer based on a MALDI-TOF mass spectrometer was used to determine the quality of produced olinscide Schip-11 (Litech, Moscow, Russia). The ratio of mass-to-charge (*m*/*z*) of olinscide Schip-11 was measured as positive ions with 3-hydroxypicolinic acid as a matrix on a LaserToFLT2 Plus device (UK) in a ratio of 2:1. The theoretical *m*/*z* ratio was calculated using ChemDraw 18.0 software (ChemDraw, Cambridge Soft, USA) and differed by no >10 units with the resulting *m*/ *z* ratio. Dilution in nuclease-free water to a required concentration was carried out on NanoDrop Lite spectrophotometer (Thermo Fisher Scientific, Waltham, MA, USA).

Concentrations of 200 ng/µL and 0.1 ng/µL of olinscide Schip-11 in nuclease-free water was applied on *P. sylvestris* leaves using hand sprayer (10 ml of water solution per m^2^ of leaves). As a control, water-treated group was used. Around 3000 insects of the 1^st^ and 2^nd^ instar were treated and their survival rates were calculated statistically. Mortality was recorded every day during 4 days for insects treated with Schip-11 in concentration 200 ng/µL and during 12 days for insects treated with Schip-11 in concentration 0.1 ng/µL. Effectiveness of olinscide Schip-11 was calculated by dividing the number of dead individuals by the total number of individuals on the shoot and multiplying by 100%, winged individuals were excluded from calculations.

### 4.3. Target gene expression

RNA isolation was performed from dead individuals. cDNA synthesis was carried out according to manufacturer’s instructions using ExtractRNA reagent (Evrogen, Moscow, Russia) and RT-PCR kit (Syntol, Moscow, Russia), respectively. RNA extraction was carried out in three replicates. For cDNA synthesis, 10 µl of RNA was taken at a concentration of 20 ng/µl. RT-PCR was performed with fluorescent dye SYBR Green. The Fast-Start SYBR Green Master Mix (Roche, Basel, Switzerland) was used according to the manufacturer’s instructions. In the SYBR Green system, DNA (2 µl) was added to the mixture with FastStart SYBR Green Master Mix kit (Roche, Basel, Switzerland) and specific primers SP_F 5’-GAC-CAG-TCG-AAC-GCA-CAT-TG-3’ and SP_R 5’-CGG-AGA-GGG-TTG-TTG-TGT-CT-3’. The expression of the target gene was evaluated on the 1^st^ and 3^rd^ day after treatment with Schip-11.

### 4.4. DNA sequencing

Total RNA was isolated from larvae using ExtractRNA kit (Evrogen, Moscow, Russia) according to manufacturer’s instructions. A MMLV RT kit was used to perform first-strand cDNA synthesis (Evrogen, Moscow, Russia), following the manufacturer’s protocols. Primers, forward 5’-GAC-CAG-TCG-AAC-GCA-CAT-TG-3’, and reverse 5’-CGG-AGA-GGG-TTG-TTG-TGT-CT-3’, were used for quantitative real-time PCR studies. PCR reactions were carried out on 2 µL of cDNA using 7 µL of FastStart SYBR Green Master Mix (Roche, Basel, Switzerland), 2 µL of ddH_2_O (Roche, Basel, Switzerland) and 0.5 µL (80 ng/µL) of each primer. DNA was first denatured for 4 min at 95 °C, then 30 cycles of 1 min of denaturation at 94 °C, 1 min of hybridization at 58 °C, and 1 min of elongation at 72 °C, followed by a final elongation step at 72 °C for 7 min. PCR products from the larvae were purified using the Cleanup S-Cap (Evrogen, Moscow, Russia) and the sequencing polymerase reaction was carried out with Big Dye Terminator v 3.1 Cycle Sequencing RR-100 (Applied Biosystems, Vilnius, Lithuania). Polymerase reactions were carried out using 2 µL of purified DNA and 2 µL of primers (12.8 ng/µL). DNA was initially denatured for 1 min at 96 °C, followed by 30 cycles of 10 s of denaturation at 96 °C, 5 s of hybridization at 50 °C, and 4 min of elongation at 60 °C. Amplicons were sequenced in both directions using the NANOPHOR-05 capillary DNA sequencer (Syntol, Moscow, Russia). DNA sequences we analyzed using ClustalW 2.0.3 program [49] and BLAST.

### 4.5. Statistical analysis

The standard error of the mean (SE) was determined and analyzed using the Student’s t-test for statistical analysis to evaluate the significance of the difference in mortality and pre-rRNA concentration between water-treated Control and experimental groups. All above-mentioned calculations were preformed using Prism 9 software (GraphPad Software Inc., Boston, USA).

## 5. Conclusions

It is believed that discoveries do not occur optimally, and it takes time for one or another significant discovery in science to find all the worthy places for its application in real life. Obviously, if P. Zamecnik and M. Stephenson in 1978 had discovered the effect of modified [8] and maybe later unmodified) antisense DNA on insect cells, then in addition to drugs, for several decades we would have had a class of insecticides based on antisense DNA [17,50]. However, history does not tolerate the subjunctive mood. Short un-modified antisense DNA as a contact insecticide was first discovered in 2008 [12] and was influenced by developments in molecular genetics, bioinformatics, and in vitro nucleic acid synthesis that have brought us into the post-genomic era in plant protection.

This article for the first time shows that low concentration of oligonucleotide insecticides (0.1 mg/L) leads to increased mortality of aphids. At a consumption rate of 200 L per hectare, the price of the required amount of oligonucleotide insecticide will be about 0.5 USD when using liquid-phase DNA synthesis. In the case of using non-optimized solid-phase DNA synthesis, the price of the required amount of oligonucleotide insecticide per hectare will be 20 USD. CUAD biotechnology opens up new frontiers for the large-scale implementation of oligonucleotide insecticides as the next generation class of insecticides in plant protection [1,2,16,22]. Oligonucleotide insecticides have a low carbon footprint [48], high safety for non-target organisms, rapid biodegradability in ecosystems, and avoidance of target-site resistance [1,16,34,36,48].

The CUAD platform is a simple and flexible biotechnology for creation of oligonucleotide insecticides [3]. Investigation of efficiency of low concentrations of oligonucleotide insecticides together with auxiliary substances (spreaders, adhesives, penetrators, etc.) will help discover the most optimal formulations for control of wide range of pests. How far we are from the point in plant protection when crop will contain only crop without traces of chemical insecticides (organic xenobiotics)? One thing is clear, we are on the way to it.

## Author Contributions

Conceptualization, V.V.O.; methodology, Y.V.P.; software, N.V.G.; validation, V.V.O.; formal analysis, Y.V.P. and N.V.G.; investigation, V.V.O.; resources, V.V.O.; data curation, Y.V.P. and N.V.G.; writing—original draft preparation, V.V.O.; Y.V.P. and N.V.G.; writing— review and editing, V.V.O.; Y.V.P. and N.V.G.; visualization, V.V.O.; Y.V.P. and N.V.G.; supervision, V.V.O.; project administration, V.V.O.; funding acquisition, V.V.O. All authors have read and agreed to the published version of the manuscript.

## Institutional Review Board Statement

Not applicable.

## Informed Consent Statement

Not applicable.

## Data Availability Statement

Not applicable.

## Acknowledgments

We thank our many colleagues, too numerous to name, for the technical advances and lively discussions that have prompted us to write this article. We apologize to the many colleagues whose work has not been cited. We are very much indebted to all anonymous reviewers and our colleagues from the Lab on DNA technologies, PCR analysis and creation of DNA insecticides (V.I. Vernadsky Crimean Federal University, Institute of Biochemical Technologies, Ecology and Pharmacy, Department of Molecular Genetics and Biotechnologies), and OLINSCIDE BIOTECH LLC. for valuable comments on our manuscript.

## Disclaimer/Publisher’s Note

The statements, opinions and data contained in all publications are solely those of the individual author(s) and contributor(s) and not of MDPI and/or the editor(s). MDPI and/or the editor(s) disclaim responsibility for any injury to people or property resulting from any ideas, methods, instructions or products referred to in the content.

